# Brief Investigation: On the rate of aneuploidy reversion in a wild yeast model

**DOI:** 10.1101/2024.09.23.614562

**Authors:** James Hose, Qi Zhang, Nathaniel P. Sharp, Audrey P. Gasch

**Affiliations:** Center for Genomic Science Innovation, University of Wisconsin-Madison, Madison, WI, 53706; Texas A&M University, School of Public Health, College Station, TX, 77843; Laboratory of Genetics, University of Wisconsin-Madison, Madison, WI, 53706

**Keywords:** Aneuploidy, mutation accumulation, chromosome nondisjunction rate

## Abstract

Aneuploidy, arising from gain or loss of chromosomes due to nondisjunction, is a special class of mutation. It can create significant phenotypic changes by altering abundance of hundreds of genes in a single event, providing material for adaptive evolution. But it can also incur large fitness costs relative to other types of mutations. Understanding mutational dynamics of aneuploidy is important for modeling its impact in nature, but aneuploidy rates are difficult to measure accurately. One challenge is that aneuploid karyotypes may revert back to euploidy, biasing forward mutation rate estimates – yet the rate of aneuploidy reversion is largely uncharacterized. Furthermore, current rate estimates are confounded because fitness differences between euploids and aneuploids are typically not accounted for in rate calculations. We developed a unique fluctuation assay in a wild-yeast model to measure the rate of extra-chromosome loss across three aneuploid chromosomes, while accounting for fitness differences between aneuploid and euploid cells. We show that incorporating fitness effects is essential to obtain accurate estimates of aneuploidy rates. Furthermore, the rate of extra-chromosome loss, separate from karyotype fitness differences, varies across chromosomes. We also measured rates in a strain lacking RNA-binding protein Ssd1, important for aneuploidy tolerance and implicated in chromosome segregation. We found no role for Ssd1 in the loss of native aneuploid chromosomes, although it did impact an engineered chromosome XV with a perturbed centromeric sequence. We discuss the impacts and challenges of modeling aneuploidy dynamics in real world situations.

**ARTICLE SUMMARY:** Errors in chromosome segregation can produce aneuploid cells with an abnormal number of chromosomes. Aneuploidy is not uncommon in wild populations of fungi and can underlie emergence of drug-resistant pathogens. But modeling the impact of aneuploidy on evolution has been challenging, because rates of aneuploidy emergence and reversion have been difficult to measure. This work used a novel fluctuation assay that incorporates euploid-aneuploid fitness differences to calculate rates of extra-chromosome loss in aneuploid *Saccharomyces cerevisiae*, across several chromosomes. The results present for the first time estimates of aneuploidy reversion (“back mutation”) rates and implications for previously measured rates of aneuploidy.

## RESULTS AND DISCUSSION

Several previous studies estimated rates of segregation errors in *Saccharomyces cerevisiae*, either in mutation-accumulation (MA) experiments to score chromosome gain (Zhu *et al*. 2014; Sharp *et al*. 2018) or marker-based fluctuation tests to study chromosome loss from diploid cells (Klein 2001; Kumaran *et al*. 2013). But a major caveat is that none of these studies to date has controlled for what may be substantial fitness differences between aneuploid mutants and their euploid progenitors. MA studies use serial population bottlenecks to limit the impact of selection, but mutations of large effect (including aneuploidy) may still be subject to selection (Lang and Murray 2008; Lang 2018; Mahilkar *et al*. 2022; Wahl and Agashe 2022), and the possibility of reversion is typically ignored. Similarly, standard fluctuation-test analysis applied to chromosome loss in diploid *S. cerevisiae* assumes that the ancestral and mutant types grow at the same rate. This is unlikely given that monosomic strains are rarely seen in nature, suggesting major fitness defects (Zhu *et al*. 2012; Peter *et al*. 2018; Scopel *et al*. 2021). Building an accurate picture of karyotype evolution therefore requires new approaches to address these sources of bias.

We devised a unique and sensitive fluctuation assay and applied novel statistical methods (Zheng 2022) to calculate rates of extra-chromosome loss. We previously generated a suite of strains with extra chromosomes in haploid oak-soil strain YPS1009 as a model wild isolate (Rojas *et al*. 2024). Aneuploids were generated using the method of (Hill and Bloom 1987), by integrating a galactose-inducible promoter next to the centromere and disrupting kinetochore attachment during division via induced transcription over the centromere. Resulting aneuploids were selected and then backcrossed to an unadulterated strain, ultimately producing haploid cells with an extra copy of an individual, native chromosome sequence.

Here, we used these strains to measure the rate of extra-chromosome loss per generation. We focused on YPS1009 strains with an extra copy of either chromosome V (Chr5), Chr13, or Chr15 because we can readily score aneuploid versus euploid colonies in these strains. We developed a strategy to select newly-born single aneuploid cells after diploid dissection, grow each cell into a population over a defined growth period, and then estimate the number of cells in the population that lost the extra chromosome for rate calculations (**Fig 1A**). We started with a diploid YPS1009 strain that was trisomic for one of the chromosomes and heterozygous for *SSD1/ssd1Δ* (discussed more below). Populations were grown for ∼20 generations from a single starting aneuploid cell. This approach enabled measurement of reversion to euploidy from aneuploid cells of both *SSD1* and *ssd1Δ* genotypes, in newly born, unaged cells grown side-by-side on the same plate.

**Figure 1.**
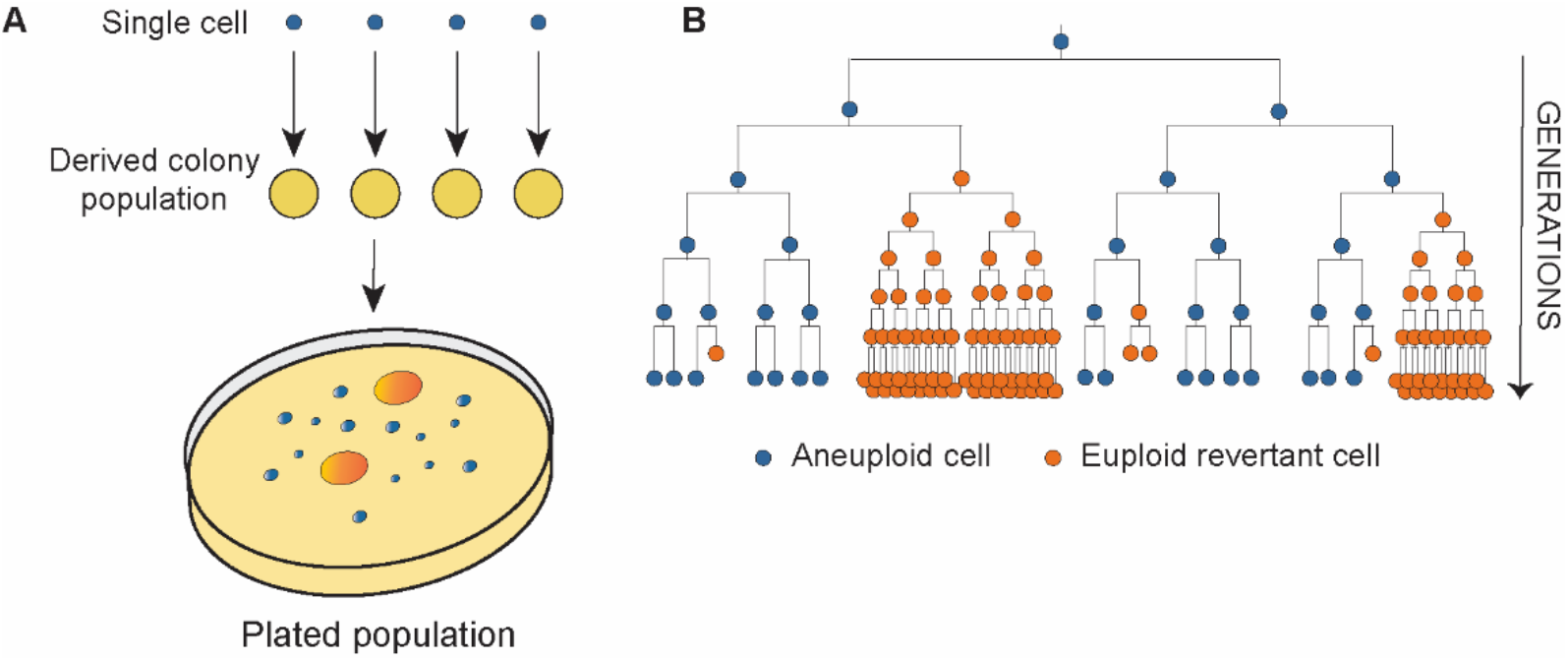
Fluctuation assay for measuring aneuploidy reversion rates. A) Procedure for measuring euploid reversion from starting aneuploidy cells. Diploid cells trisomic for the chromosome of interest are sporulated, and the meiotic products produce single cells of euploid and aneuploid genotypes, with and without *SSD1*. Aneuploid cells are allowed to grow into a colony for ∼20 generations, during which time some cells stochastically lose one copy of the extra chromosome during cell division. After a defined growth period, each colony is scraped and cells in the colony “population” are isolated and replated. Single cells that are euploid at the time of replating grow at euploid rates (defined carefully in control strains and validated with quantitative PCR) to produce relatively large secondary colonies. In contrast, cells that remain aneuploid at the time of replating grow more slowly into colonies of distinctly smaller size and with unique morphology (see Methods). The number of euploid revertants across populations was used to model rates. B) Stochastic loss of the extra chromosome, coupled with significant fitness differences in euploid cells, can lead to jackpot numbers of euploids in the population that must be accounted for in the modeling.

The stochastic nature of chromosome loss results in variation in the number of euploid cells across populations, depending on the number and timing of chromosome loss events (**Fig. 1B** and (Luria and Delbruck 1943)). Mutation rates can be estimated based on the distribution of mutant counts across populations (Luria and Delbruck 1943; Lea and Coulson 1949; Lang 2018). However, in our case euploid cells also grow much faster than aneuploids, which we hypothesized would bias rate estimates. We therefore applied a modified algorithm specifically designed for this case (Zheng 2022). An important parameter in our calculation is the relative fitness of euploid versus aneuploid cells, sensitively measured based on aneuploid and euploid colony growth rates (see Methods). We therefore calculated relative fitness *w* of euploids versus aneuploids based on their relative doubling times measured in our system. The resulting analysis produced rates of chromosome loss per generation along with confidence intervals (CIs) that enable direct comparison across strains and chromosomes (Zheng 2015).

### Controlling for fitness differences significantly impacts mutation rate estimates

We estimated rates of extra-chromosome loss for cells carrying specific chromosome duplications, starting in *SSD1+* cells. Accounting for fitness differences revealed that rates of loss ranged from 8.3 × 10^−5^ for Chr15 to 6.8 × 10^−4^ and 5 × 10^−4^ for Chr5 and Chr13, respectively (**Fig. 2A**). To test how neglecting fitness could bias rates, we re-calculated rates without accounting for fitness, setting relative fitness *w* to 1.0. In all cases, this resulted in an over-estimation of extra-chromosome loss rates (**Fig. 2A**). In general, this suggests that rates of mutation towards less-fit karyotypes (disomy in haploids, monosomy or trisomy in diploids) are also likely to be underestimated (see *Perspective* below).

**Figure 2.**
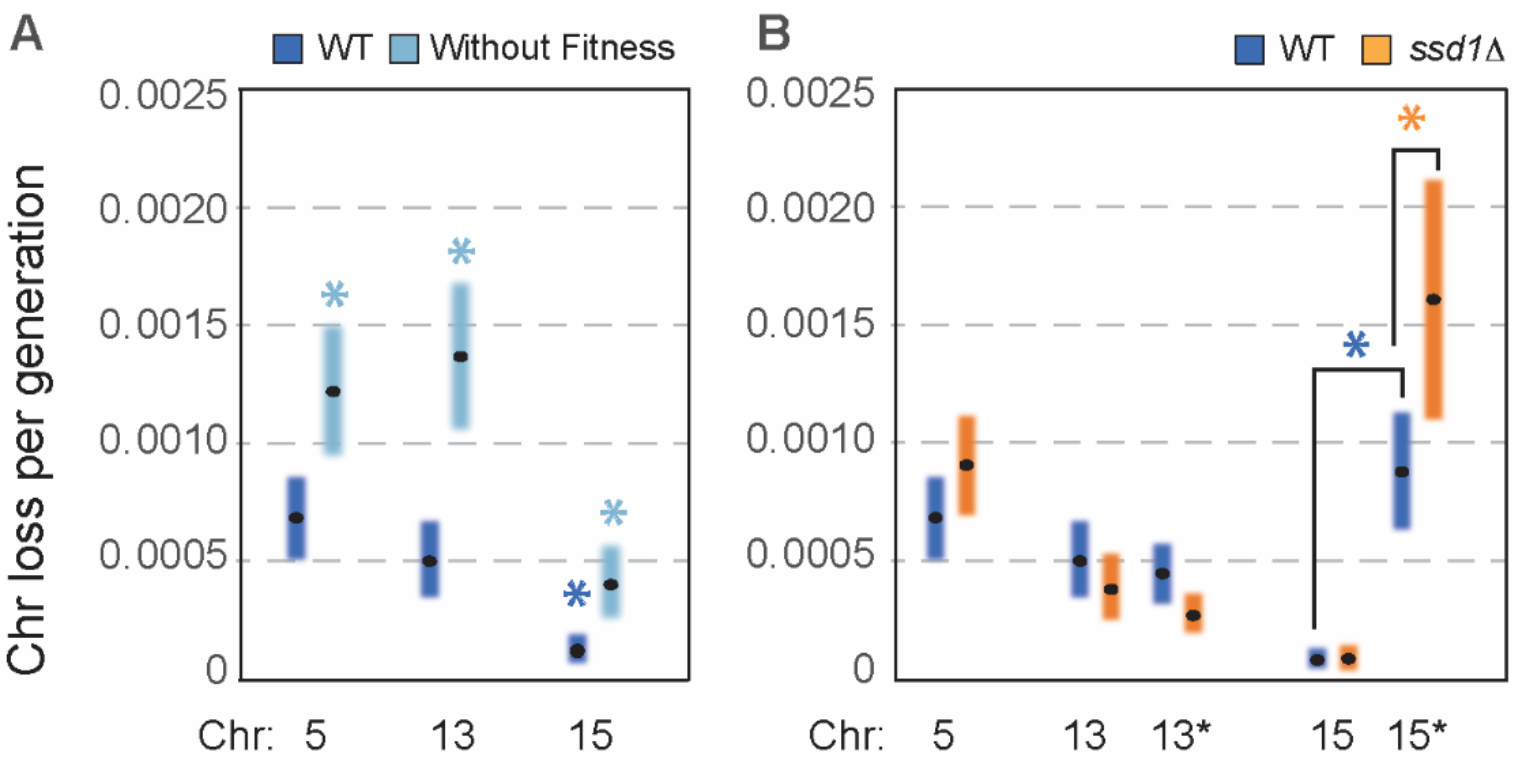
Rates of extra-chromosome loss. A) The rate of loss of extra Chr5, Chr13, and Chr15 (dark blue) is shown by each point in the figure, surrounded by the 95% confidence interval (CI), for wild type cells (dark blue). The rate of extra-Chr15 loss is significantly lower than the other chromosomes (dark blue asterisk, p<0.05, comparing 83% CI (Zheng 2015)). Rates calculated when fitness was ignored are shown in light blue. In all cases, the rate was significantly different when fitness was ignored in the calculation (asterisk, p<0.05). B) Rates of extra-chromosome loss as shown in A for wild-type cells (reprinted from A for direct comparison) and cells lacking *SSD1*, for native chromosome sequences or in chromosomes that retain the centromere-proximal cassette (Chr13*, Chr15*). The loss of Chr15* was over 10-fold higher than the native Chr15 chromosome in wild-type cells (dark blue asterisk, p<0.05). Loss of the marked chromosome was an additional 1.8-fold higher in *ssd1Δ* cells compared to wild-type cells with the same Chr15* chromosome (orange asterisk, p<0.05), showing that Ssd1 influences that rate of Chr15* loss when the cassette is retained on the chromosome.

### Extra chromosomes are lost at different rates

Different aneuploidy karyotypes impose different fitness costs, depending on chromosome size, gene content, and other factors (Rojas *et al*. 2024). Previous studies suggested that different chromosomes may be lost at different rates in diploid cells; however, an alternative explanation is that unaccounted-for differences in the relative fitness of aneuploids bias the results. Thus, it has been difficult to assess if chromosomes are lost at different inherent rates. Here we show by accounting for fitness differences that extra chromosomes are indeed lost at different rates. Whereas the rate of loss of aneuploid Chr5 and Chr13 were not statistically different (p>0.05 based on overlapping 83% CIs), the rate of extra-Chr15 loss was 6 to 8-fold lower (**Fig. 2A**).

The mechanism of chromosome-specific loss rates could be explained by several factors. Kumaran et al. (2013) argued that monosomy rates are inversely proportionate to chromosome size; this agrees with results from artificial chromosomes in which small chromosomes are lost at a higher rate (Hieter *et al*. 1985; Murray *et al*. 1986). However, because aneuploidy fitness costs are highly correlated with chromosome size (Torres *et al*. 2008; Gilchrist and Stelkens 2019; Scopel *et al*. 2021; Rojas *et al*. 2024), an alternative explanation is that loss of larger chromosomes to produce monosomy incurs a larger fitness cost, minimizing their detection in diploid cells and producing proportionately greater bias for larger chromosomes.

By controlling for fitness differences, our study offered an opportunity to investigate this question. We found that over this small set of chromosomes, the rate of extra-chromosome loss did not correlate with chromosome size. Based on the sequenced YPS1009 genome (Rojas *et al*. 2024), Chr13 is 1.6 times larger than Chr5, yet the rate of loss of these extra chromosomes was not significantly different. In contrast, Chr15 is comparable size to Chr13 (1.16 times its size) – yet it is lost at 6-fold lower rate than Chr13. Thus, although we interrogate only a small number of chromosomes here, their loss rate is not tightly coupled with chromosome size, at least not in this strain background. This conclusion parallels results from MA lines (Zhu *et al*. 2012). An alternative hypothesis is that chromosome-specific rates of loss are influenced by other chromosome features, including genes encoded on that chromosome, differences in centromeric sequence, or variation in local context that affects centromere function (Panzeri *et al*. 1985; Sears *et al*. 1995; Bechert *et al*. 1999; Bensasson 2011; Ohkuni and Kitagawa 2012; Bloom and Costanzo 2017). In fact, the efficiency of plasmid maintenance is influenced by the identity of the cloned centromere on the plasmid (Kumaran *et al*. 2013). Thus, chromosome-specific features likely influence the rate of extra-chromosome loss when fitness is accounted for.

### Ssd1 does not affect native chromosome loss but influences loss of modified Chr15

The RNA binding protein Ssd1 is critically important for tolerating extra chromosomes (Hose *et al*. 2020; Dutcher *et al*. 2024; Rojas *et al*. 2024), although the mechanisms through which it enables aneuploidy tolerance remain incompletely understood. Ssd1 has been implicated in plasmid maintenance and binds several mRNAs encoding proteins important for chromosome segregation (including several kinetochore and spindle-pole body proteins), suggesting that it may be important for the stability of the extra chromosomes (Uesono *et al*. 1994; Hose *et al*. 2020). Indeed, *ssd1-* aneuploid cultures readily revert to a euploid population – however whether this is because Ssd1 plays a role in chromosome maintenance or is simply due to the extreme competitive disadvantage of *ssd1-* aneuploids versus the euploid remained unclear.

We therefore used our sensitive assay to test if the rate of loss of the extra-Chr5, Chr13, and Chr15 was higher in cells lacking *SSD1*. For all of the chromosomes studied, the rate of loss was indistinguishable for *ssd1Δ* aneuploids versus the wild type (**Fig. 2B**). Thus, to the extent that we can measure here, Ssd1 does not play a major role in the stability of these aneuploid chromosomes. Instead, the propensity to lose aneuploidy in *ssd1Δ* cultures is driven by the severe fitness defect of *ssd1Δ* cells, coupled with the relatively high rate of stochastic chromosome loss.

However, during strain construction we noticed that cells harboring an extra Chr15 that retained the galactose-inducible, centromere (CEN)-proximal cassette used to induce segregation errors reverted more frequently to large colonies on a plate. This suggested that the CEN-proximal cassette could destabilize chromosomes, even in the absence of galactose inducer. We therefore measured rates of loss of Chr13 and Chr15 that retained the cassette (Chr13* and Chr15*). Chr13* with the cassette was lost at an indistinguishable rate compared to the native Chr13 (**Fig. 2B**). However, the CEN cassette destabilized Chr15, since Chr15* was lost at a 10-fold higher rate than the native chromosome without the cassette. This was surprising, since there was no inducer present in the media. Remarkably, Chr15* was lost at an even higher rate in *ssd1Δ* cells compared to wild-type (1.7 × 10^−3^ in *ssd1Δ* Chr15* cells versus 8.7 × 10^−4^ in wild-type Chr15* cells, p<0.05). Thus, chromosome stability was differentially affected by the CEN-proximal cassette, depending on the chromosome and *SSD1* status.

We designed several alternate cassettes and measured rates of chromosome loss to test models (**Fig. 3A**). Rates were indistinguishable from the unmarked, native Chr15 chromosome when GFP and/or the hygromycin-resistance gene were integrated next to the centromere (**Fig. 3A**, constructs b and d), indicating that DNA insertion at that locus does not have a major impact on segregation. However, cassettes containing the GAL1-10 promoter facing the centromere (**Fig. 3A**, constructs a and c) substantially increased loss rates, in a way that was independent of the *GAL4* transcription factor (**Fig 3B**). Recent evidence shows that the *GAL1-10* promoter is subject to leaky transcription in the absence of an antisense noncoding transcript emerging from the downstream *GAL10* gene, even in the absence of galactose (Lenstra *et al*. 2015). We propose that leaky *GAL1-10* expression is amplified in a Chr15- and Ssd1-dependent manner. One possibility is that the effect is influenced by amplified genes on Chr15 that include *GAL11*, encoding a subunit of the RNA Pol II mediator complex to which Ssd1 has been shown to bind (Phatnani *et al*. 2004). Future work will be required to test this hypothesis; nonetheless, we conclude that the role of Ssd1 in tolerating extra chromosomes is unlikely to be through a role in maintenance of native extra-chromosome sequences.

**Figure 3.**
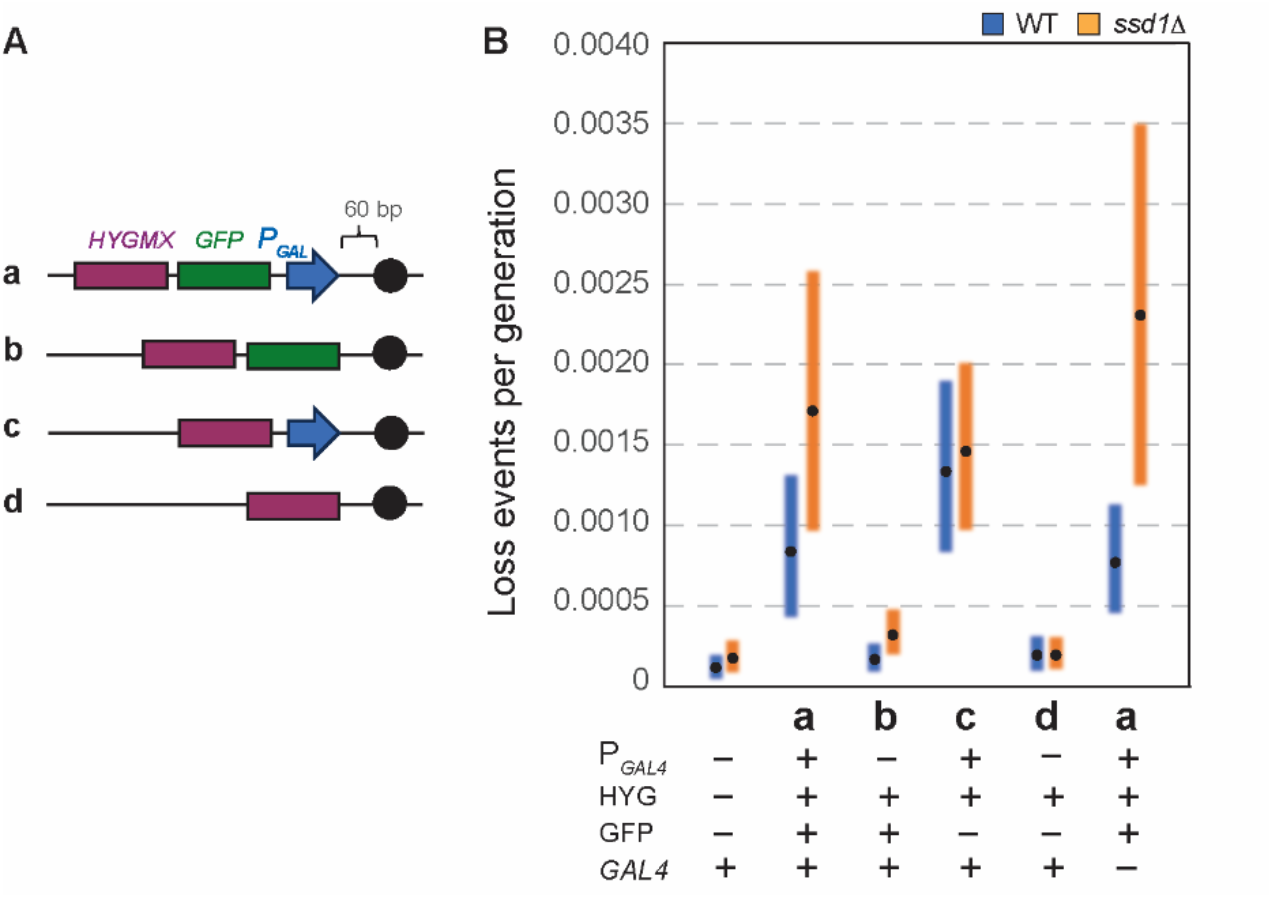
The GAL promoter influences Chr15* stability independent of Gal4 inducer. A) Diagram representing cassettes tested, see main text and Methods. The original cassette used during strain construction is (a). B) Rates of extra-Chr15 loss in wild-type (blue) and *ssd1Δ* (orange) cells depending on the cassette shown in A.

## PERSPECTIVE

Our results have implications for the interpretation of aneuploidy rate estimates from previous experiments. Fluctuation tests will produce biased results if fitness differences are not accounted for, and we show that this bias can be substantial in the case of aneuploidy. The direction of bias will depend on whether the ancestral cell type has higher or lower fitness than the “mutant” type; in general, we expect that failing to account for fitness will result in over-estimation of the rate of reversion to a euploid state, and under-estimation of the rate of aneuploidy appearance. For example, reported rates of chromosome loss in diploid cells for the same chromosomes we studied here are ∼3 × 10^−6^ for Chr5, <2 × 10^−8^ for Chr13, and ∼2 × 10^−7^ for Chr15 (Klein 2001; Kumaran *et al*. 2013). On its face, this comparison suggests that chromosome loss in diploids is 3-5 orders of magnitude rarer than the loss of an extra chromosome in haploids. However an important consideration is that unaccounted-for reduced fitness in monosomic diploids generates a downward estimation bias.

Similarly, MA experiments may under-estimate the rate that aneuploidy arises, if the deleterious effects of mutant karyotypes are stronger than neutral genetic drift under repeated bottlenecking. Furthermore, MA studies typically do not account for the potential for reversion (back mutation), which is unlikely to affect point mutation due to the low probability of repeated mutations at the same site. However, this is not the case for aneuploidy, where rates of mutation are several orders of magnitude higher than point mutation rates (Zhu *et al*. 2012; Sharp *et al*. 2018). The downward bias due to back mutation will depend on the relative forward and reverse mutation rates. A study of 106 haploid yeast MA lines for ∼1490 generations observed no instances of chromosome gains, suggesting a mutation rate to disomy no greater than ∼1.5 × 10^−6^ (95% confidence limit on a per-chromosome basis; (Sharp *et al*. 2018) see also (Serero *et al*. 2014). In concert with our results, this implies a much lower rate of forward mutation (chromosome duplication) in haploids than back mutation (extra-chromosome loss). A higher rate of back mutation could emerge if aneuploidy creates chromosomal instability (Sheltzer *et al*. 2011; Zhu *et al*. 2012) – but this result could also reflect biases in previous estimates of chromosome gain. Although strong selection against disomy could prevent its accumulation in MA, disomic mutants have been observed in mutator-strain MA studies (Serero *et al*. 2014), indicating that disomy is at least partially detectable under MA. But MA experiments may also miss aneuploidy events if they revert at an appreciable rate – our results highlight the plausibility of this scenario. Diploid MA lines accumulate aneuploidy events at an apparent average rate of ∼6 × 10^−6^ per chromosome, but with variability among chromosomes, a deficit of losses (monosomy) relative to gains (trisomy), and evidence for reversion events (Zhu et al. 2014, Sharp et al. 2018). At present, it is unclear to what extent these patterns reflect genuine variability in forward and reverse mutation rates versus selection on alternative karyotypes. There is a clear need for more data on rates of aneuploidy formation and loss that accounts for potential sources of bias.

An additional challenge modeling aneuploidy dynamics is that fitness costs of aneuploidy are highly dependent on environmental conditions but also genetic background (Zhu *et al*. 2016; Gilchrist and Stelkens 2019; Hose *et al*. 2020; Scopel *et al*. 2021; Li and Zhu 2022; POMPEI AND COSENTINO LAGOMARSINO 2023). Fitness costs can even vary with growth phase, since several studies show that chromosome amplification in yeast leads to shortened life span, separable from growth rates of young cultures (Sunshine *et al*. 2016; Escalante *et al*. 2024). Thus, how cells adapt to fluctuating environments especially when fitness costs fluctuate is an area of active study (Abreu *et al*. 2024). In addition to fitness effects, mechanisms affecting chromosome loss can vary across strains, producing strain-by-chromosome effects on chromosome maintenance. Indeed, centromeric sequences where kinetochores attach evolve at a fast rate (Bensasson 2011; Roach *et al*. 2012; Logsdon *et al*. 2024). These features complicate modeling of aneuploidy dynamics in real-world situations. Nonetheless, continued focus on defining fitness effects, rates of forward and back mutation for aneuploidy states, and mechanisms behind their variation will continue to enhance understanding of the impact of aneuploidy on evolution.

## METHODS

### Strains and growth conditions

Strains used are listed in Table 1A. Chromosome loss assays were initiated with freshly created diploid YPS1009 cells lacking the *HO* endonuclease, trisomic for the respective chromosome, and heterozygous for *SSD1/ssd1Δ* to enable scoring multiple genotypes in the same assay by tracking single cells after dissection. Matings to generate those diploids are listed in Table 1B. Cell genotypes were determined *post hoc* after sporulation (described below). Strain AGY2047 was generated by replacing the *URA3* gene on Chr5 with *Schizosaccharomyces pombe HIS3* gene; this strain was used to generate diploid strains with *URA3* and *HIS3* marked copies of Chr5 (Table 1B). Unless otherwise noted, strains were grown in YPD rich medium (1% yeast extract, 2% peptone, 2% dextrose). Maintenance of the aneuploid chromosome in starting strains was verified periodically by qPCR of select gene(s) on and off the amplified chromosome, to ensure ∼2X copy of the expected amplified genes.

### Chromosome Loss Assay

We scored the number of cells in a population that lost the extra chromosome in question as outlined in Fig. 1. We began with diploid *ho-/ho-* YPS1009 cells with three copies of either Chr5, Chr13 or Chr15 and heterozygous for *SSD1* (see Table 1). Each diploid strain was sporulated on solid medium with 1% potassium acetate for two days, and tetrads were dissected on solid YPD medium at defined spacing, producing haploid *SSD1+* and *ssd1Δ* spores with 1 (euploid) or 2 (aneuploid) copies of the chromosome in question. In this strain background, viability of euploid/aneuploid segregants is nearly 100%, and tetrads routinely show 2:2 segregation of euploid:aneuploid cells as expected, determined by qPCR and/or colony size. The dissection plate was incubated for 32-48 h, depending on the aneuploid dissected, with the aim of achieving ∼20 generations of growth on the plate. A benefit of this procedure is that genotypes being compared are grown side-by-side, from cells born anew after meiosis, thereby minimizing effects of cellular age, differential nutrient availability, or growth times.

Individual colonies (defined as the “population” in Fig. 1) that arose from each single starting cell were scraped from the plate and resuspended in liquid YPD medium at a density of ∼200 cells / µL. The genotype of each starting cell was predicted based on colony size at the time of selection (see more below) and confirmed *post hoc* based on segregation of drug resistance or prototrophy (hygromycin resistance for chromosomes with the *HYGMX*-marked cassette, uracil or histidine prototrophy for YPS1009_Chr5 cells, or G418 resistance for *ssd1Δ::KANMX* marked cells). Cells in each suspension were counted on a Guava flow cytometer (Milipore, Billerica, MA) to produce *N*_*t*_, the number of cells in each population. *N*_*t*_ was adjusted to estimate viable cells for *ssd1Δ* cells with extra Chr13 or Chr15, which showed reduced viability at 83% and 80% based on control platings. An average of 15 populations (each emerging from a single cell) were interrogated per genotype, generated across at least 3 different days (with the exception of the *gal4Δ* strains generated on 2 separate days). In each experiment, the doubling time of euploid and aneuploid cells was estimated based on the average number of cells in each population, for colonies grown side-by-side on the same dissection plate. Relative fitness (w) of each aneuploid versus euploid revertants was taken as the ratio of doubling times for aneuploid versus euploid cells in that experiment.

To quantify the number of euploid revertants (y_k_) in each sample of plated cells, a portion of each population (aiming for ∼5,000 cells) was spread evenly onto ten 150 mm petri dish with solid YPD medium and grown for 48 h. The fraction of viable cells from the population that were scored defines the plating efficiency (ε) for each population. Euploid revertants in each population were determined by several methods, maximizing accuracy in the calls. YPS1009 disomic for Chr13 or Chr15 can be readily distinguished from euploid colonies on a plate due to their proliferative defects producing smaller colonies (Hose *et al*. 2020) along with unique colony morphology (wherein colonies that start out aneuploid develop ruffled edges due to the accumulation of faster-growing euploid colonies along the periphery). Both colony size and morphology show a clear dichotomy for aneuploids and euploids that we verified extensively by qPCR, genomic sequencing, and post-analysis assessment of aneuploidy maintenance through differentially marked aneuploid chromosomes (Hose *et al*. 2020; Dutcher *et al*. 2024; Rojas *et al*. 2024). Petite colonies that have lost respiratory capacity are relatively few in this genetic background and readily apparent based on colony color, and thus they were not scored in the analysis. Colony-size difference between YPS1009 disomic for Chr5 (YPS1009_Chr5) versus euploid cells is more subtle but still distinguishable. Therefore, potential euploid-revertant colonies scored based on size were also assessed *post hoc* for loss of one of two auxotrophic markers (*HIS3* or *URA3*) present on either copy of Chr5. Cells were streaked onto minimal media plates without uracil and/or histidine to verify that one or other marker had been lost, consistent with loss of that chromosome. Cells were also confirmed by colony morphology described above.

Thus, colonies emerging from YPS1009_Chr5 aneuploids were scored as euploid revertants if they were larger, lost one of the auxotrophic markers, and produced smooth, round colonies that lacked the aneuploid colony morphology.

## Modeling of chromosome loss rates

Because we relied on a modified Luria-Delbrück fluctuation assay protocol (see Fig. 1 and (Luria and Delbruck 1943)), our mathematical model for inference about aneuploidy loss rates was based on a modified version of the classic Lea-Coulson distribution (Lea and Coulson 1949). Specifically, the Lea-Coulson mutant distribution does not incorporate two important features in our system: differential fitness of euploid versus aneuploid strains and low plating efficiencies that score only a fraction of each population. To our knowledge, there existed no practical algorithms for computing the mutant distribution under this modified Lea-Coulson model. Therefore, we applied an algorithm inspired by the experiment in this investigation (Zheng 2022). This algorithm enabled maximum likelihood estimates of the aneuploidy reversion rates *μ* and their corresponding confidence intervals (CIs). Parameters in the model included differential fitness *w*, the number of viable cells in each population at the time of plating, *N*_*t*_, the plating efficiency, ε, and the number of observed euploid revertants *y*_*t*_ in each plated aliquot of the population. Our method also allowed for a statistical procedure to test rate differences by checking overlapping of 83% CIs, which corresponds to p = 0.05 (Zheng 2015). Python code to perform the analyses is available at https://eeeeeric.com/rSalvador/. Input data to the algorithm is provided in Table 2. Estimated rates and confidence intervals are provided in Table 3.

## TABLE LEGENDS

**Table 1: Strains used in this study.** A) Strains and their genotypes and sources. B) Crosses used in the analysis. Diploid strains were generated freshly by mating designated strains, and then sporulated as described to produce haploid genotypes of interest (see Methods). The centromere-proximal cassettes reflect the full cassette (cen-a, see Rojas et al.) or derivatives cen-b, cen-c, cen-d as shown in Figure 3A.

**Table 2: Raw data used in calculations.** Measured values for different experiments (annotated by Experiment Name ID) including indicated parameters in the model such as population size *N*_*t*_, plating efficiency ε, number of euploid revertants counted *y*_*t*_, and calculated fitness difference *w*.

**Table 3: Calculated rates and confidence intervals (CI).**

